# Neuronal population activity dynamics reveal a low-dimensional signature of operant learning

**DOI:** 10.1101/2021.12.06.471434

**Authors:** Renan M. Costa, Douglas A. Baxter, John H. Byrne

## Abstract

Learning engages a high-dimensional neuronal population space spanning multiple brain regions. We identified a low-dimensional signature associated with operant conditioning, a ubiquitous form of learning in which animals learn from the consequences of behavior. Using single-neuron resolution voltage imaging, we identified two low-dimensional motor modules in the neuronal population underlying *Aplysia* feeding. Our findings point to a temporal shift in module recruitment as the primary signature of operant learning.

## Main Text

Understanding neuronal population activity spanning multiple brain regions has emerged as central to understanding brain functions. The strategy of building low-dimensional representations of high-dimensional population activity offers insights into the generation of behavior^1–3^, learning^1,4,5^, and other functions^6,7^. However, monitoring activity in a substantial proportion of the population is only feasible in a few systems (e.g., ^1,3^). Therefore, it remains unknown whether it is possible to identify a low-dimensional “*learning signature*” associated with operant conditioning (OC), a ubiquitous form of learning in which an animal learns from the consequences of its behavior.

We addressed this question by leveraging advantages of the well characterized feeding behavior of *Aplysia*, which exhibits OC^8,9^. The feeding circuit in the buccal ganglia, including some loci of nonsynaptic and synaptic plasticity engaged by OC, has been characterized^10-13^. Isolated ganglia continue to generate buccal motor patterns (BMPs) (Fig. 1), and activity in a substantial proportion of the BMP-generating circuit can be monitored using voltage-sensitive dye (VSD) imaging^14,15^. We combined VSD imaging with an *in vitro* analogue of OC^16^ to examine reconfiguration of population activity induced by learning.

**Fig. 1.**
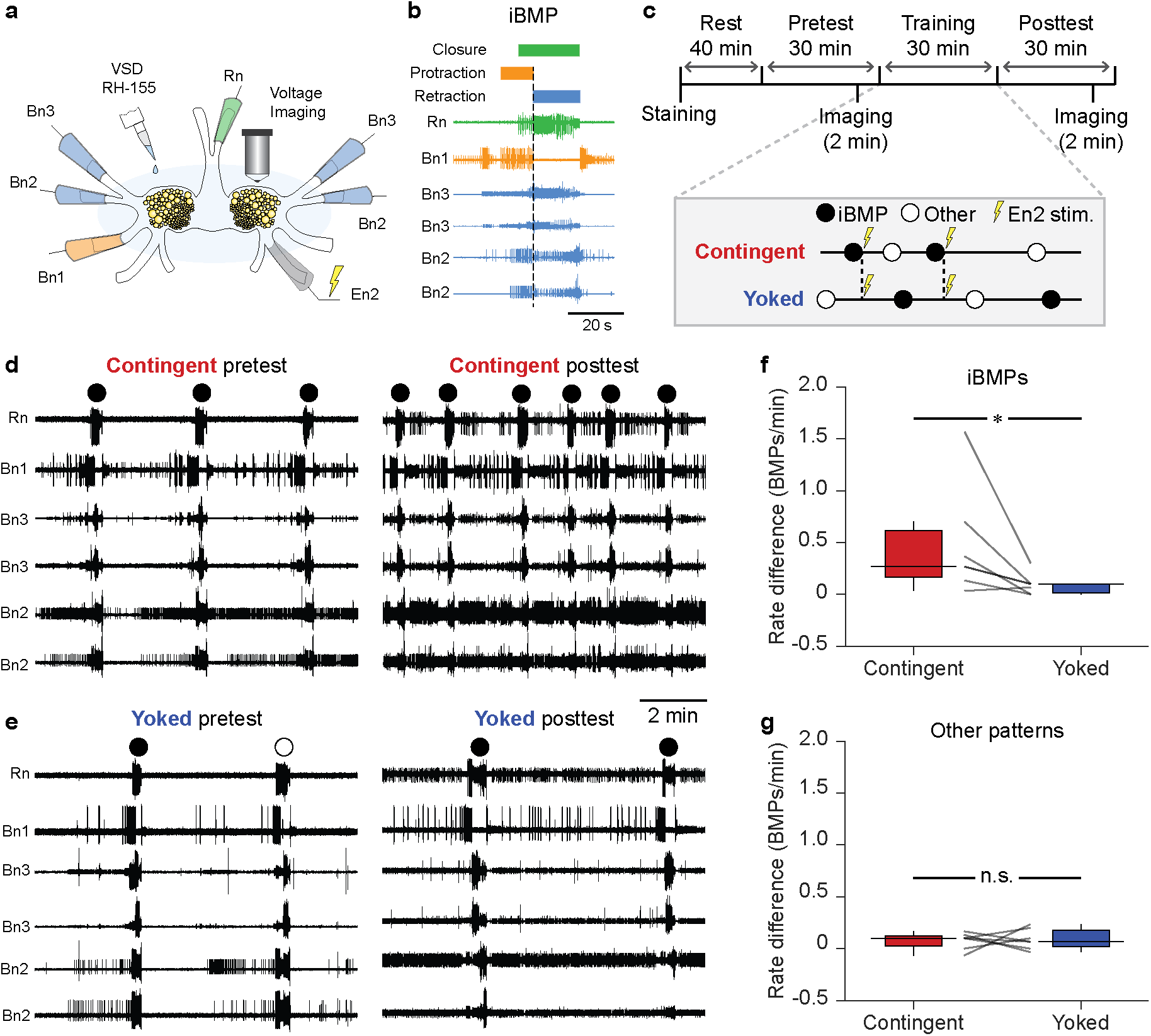
*In vitro* analogue of operant conditioning. **a**, Schematic of isolated ganglia for simultaneous extracellular electrophysiology and VSD imaging. **b**, Example BMP. **c**, Timeline and training paradigm. **d, e**, Examples of nerve activity before and after contingent (d) or yoked training (e). **f**, Contingently trained preparations had a greater increase in the rate of iBMPs compared to yoked preparations (Wilcoxon signed-rank test, P = 0.03, W = −26, N = 7; medians: contingent = 0.27, yoked = 0.10 BMPs/min). **g**, No significant difference was observed for other patterns (P = 0.94, W = −2, N = 7; medians: contingent = 0.10, yoked = 0.07 BMPs/min). BMP rates were pretest-subtracted. Individual datums represent preparations. Box plots show median (line), quartiles (box) and range (whiskers).

We used extracellular suction electrodes in buccal nerves 1–3 (Bn1–3) and the radula nerve (Rn) (Fig. 1a–b) to record BMPs before and after OC (Fig. 1c–e). Direct stimulation of the dopamine-rich esophageal nerve 2 (En2), which acts as a reward, was made contingent upon ingestion-like BMPs (iBMPs). Each contingent preparation was paired with a yoked control that received identical stimuli irrespective of its own BMP occurrences (Fig. 1c). Consistent with previous studies, conditioning specifically increased the rate of iBMPs compared to yoked controls, but did not change the rate of other, non-iBMP patterns (Fig. 1f–g).

To gain insight into the effects of OC on high-dimensional population dynamics, OC was combined with population-wide, single-neuron resolution VSD imaging (Fig. 2). We monitored ∼100 neurons per preparation. About one-third exhibited activity (Fig. 2a–c). Many neurons displayed correlated phasic firing, similar to previous studies^14,15^.

**Fig. 2.**
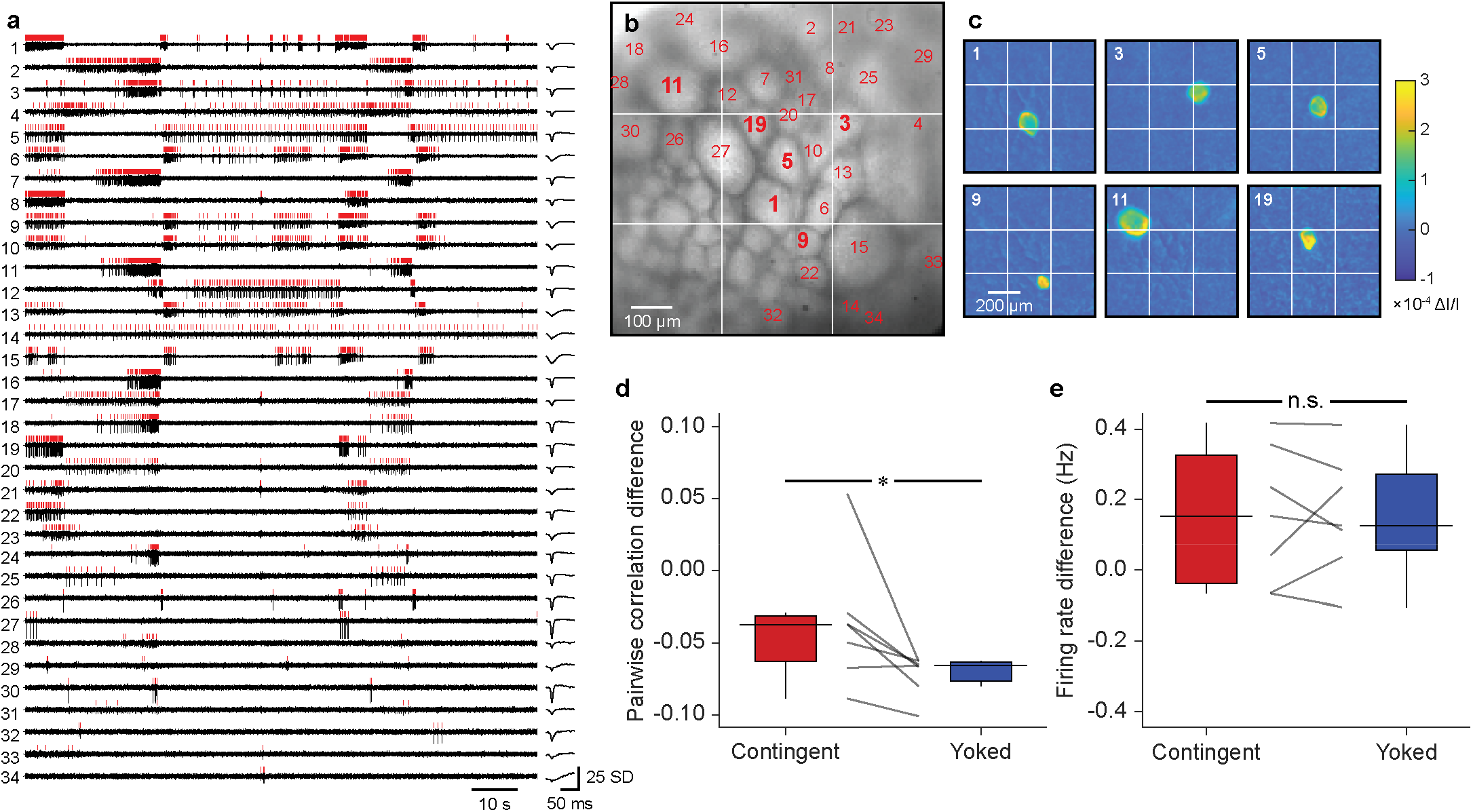
VSD imaging captures changes in primary features of population activity. **a**, Z-scored light intensities of neurons (left, black) and average waveform of detected action potentials (right). Red vertical lines indicate spike times. Neurons that had at least 5 spikes during the 120-s recording period are displayed (34 out of 92 neurons). **b**, Imaged ganglion. Numbers correspond to neuron IDs of the cells in panels a and c. **c**, Localized changes in light intensity during action potentials. Each panel displays the average change in intensity from baseline to peak across all detected spikes in the cell. Neuron ID shown on top left of each panel. **d**, Contingent training led to higher pairwise correlation across the population than yoked training (P = 0.03, W = −26, N = 7; medians: contingent = −0.04, yoked = −0.07). **e**, No significant difference was observed in overall firing rate across the population (P = 0.81, W = −4, N = 7; medians: contingent = 0.15, yoked = 0.13). Data were pretest subtracted. Individual datums represent preparations. Box plots show median (line), quartiles (box) and range (whiskers).

As a first step toward characterizing population-level features of operant reward learning, we examined overall changes in pairwise firing correlation and firing rate. We computed the mean firing correlation across all pairs of neurons within each preparation, and asked whether this correlation matrix differed between contingently-trained and yoked-control preparations. Contingently trained neuronal populations were significantly more correlated than those with yoked training (Fig. 2d) without a clear difference in overall firing rate (Fig. 2e). Thus, a distinctive feature of operant reward learning was enhanced coincident firing among units, rather than altered level of activity.

The high dimensionality of the neuronal population space hinders further intuitive interpretation. Consequently, we asked whether there is a low-dimensional subspace that captures most of the dynamics, using the dimensionality reduction approach non-negative matrix factorization (NNMF)^17^. For each preparation, NNMF extracted two modules from the high-dimensional population activity (Fig. 3a, Extended Data Fig. 1c). Each module is defined by: 1) a set of weights or contributions from each neuron, which represents the subset of neurons participating in the module, and 2) a magnitude or level of recruitment of the module at each point in time. Neuron participation in the modules was sparse and largely non-overlapping. Together, these modules explained 74.9 ± 1.8% of the power in the data, suggesting they captured most of the population dynamics while reducing the dimensionality to two.

**Fig. 3.**
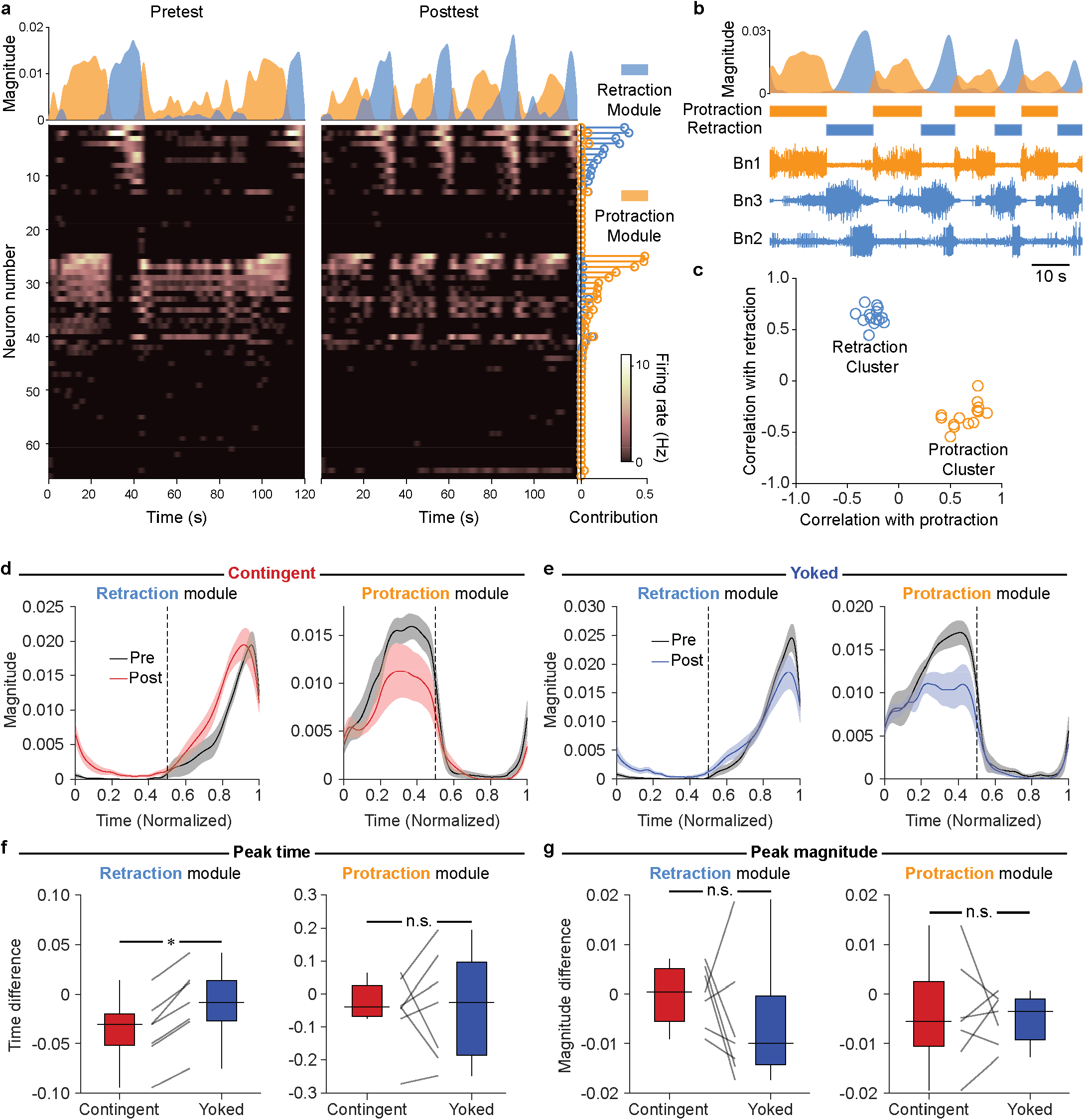
Contingent training advances recruitment of the retraction, but not the protraction, motor module. **a**, Example NNMF of population activity into two modules. Area plots (top) show module recruitment over two 120-s recording periods. Stem plots (right) indicate the module contributions by each neuron. Heat maps (bottom) show activity. Neurons are sorted by the difference between contributions to each module. **b**, Example of module recruitment (top) during a sequence of BMPs. BMP phases (middle) were labeled based on nerve recordings (bottom). **c**, Correlation between module recruitment and BMP phases across animals. Modules formed two clear clusters—one correlated with retraction, and the other with protraction. **d**, Module recruitment before (black) and after (red) contingent training, shown over the course of a BMP normalized for phase duration. Dashed lines represent transition between protraction and retraction phases. Lines and shading represent mean and standard error. **e**, Same as panel d, but for yoked preparations. **f**, Temporal shift following training. Recruitment of the retraction module was significantly advanced (P = 0.02, W = 28, N = 7; medians: contingent = −0.03, yoked = −0.01). No significant difference was observed for the protraction module (P = 0.81, N = 7). **g**, Change in magnitude of peak module recruitment was not statistically significant for either retraction (P = 0.30, N = 7) or protraction (P = 0.81, N = 7) modules. Individual datums represent preparations. Box plots show median (line), quartiles (box) and range (whiskers).

Given that NNMF can decompose behavior^18^, we asked whether the modules corresponded to specific aspects of BMPs. BMPs consist of two main phases— protraction followed by retraction. Although neither phase timing nor nerve activity were used to obtain the modules, module recruitment overlapped with either protraction or retraction (Fig. 3b). Indeed, each module across all preparations was correlated with the timing of one phase and anti-correlated with the other, forming two clusters (Fig. 3c). Modules were also highly consistent across preparations (Extended Data Fig. 1). These findings suggested the observed population dynamics reflected recruitment of neurons participating in a retraction and a protraction motor module for each animal.

Next, we asked whether a low-dimensional signature for operant reward learning was apparent in these motor modules by examining recruitment during BMPs before and after contingent or yoked training (Fig. 3d–e). The peak recruitment of the retraction module was significantly advanced in contingent preparations compared to yoked controls. In contrast, no significant difference in timing was observed for the protraction module (Fig. 3f). Interestingly, there was no difference between groups in the peak recruitment magnitude of either the retraction or protraction modules (Fig. 3g). These findings suggest the most prominent feature of OC was a change in the relative timing, but not the level, of population activity. Furthermore, that change was specific to a particular subset of neurons within the population (those participating in the retraction module).

Many neurons in the ganglia have been uniquely identified and characterized in previous studies (e.g., ^11,19^). Thus, one can ask which neurons are consistently part of the cells modulated by OC across animals. Population-wide imaging does not provide information on features typically used to identify neurons, such as synaptic and intrinsic properties. We therefore examined the overall localization of neurons contributing to the retraction and protraction modules, and compared this distribution to previously characterized neurons (Extended Data Fig. 2). Neurons that strongly contributed to the retraction module were typically localized to the top-left quadrant, whereas neurons contributing to the protraction module tended to be localized more centrally and broadly (Extended Data Fig. 2a). Neurons contributing to both modules stretched from medial to lateral, but neurons contributing to the retraction module were more ventral (Extended Data Fig. 2b). Localization of both modules was consistent with previously described neurons known to be active during either retraction or protraction (Extended Data Fig. 2c). In addition, the region with the most consistent and strong contributions to the retraction module appeared to coincide with the ventral region where several retraction motor neurons are located (e.g., B3, B6, B7, B9, B10, B39, B43, B44). Determining the extent to which biophysical changes in these neurons are responsible for the shift will require extensive electrophysiological studies focusing on them and on cells that could affect their activity, but that were likely not part of the retraction module due to their localization (e.g., B51, B64). Why is activity modified specifically in the retraction module? All outcome-based learning systems must address the credit assignment problem. That is, following an outcome such as a reward, which neurons or synapses should be modified and to what extent? Temporal contiguity is a primary mechanism of credit assignment^20–23^. In our training, reward immediately followed retraction termination. Furthermore, the retraction module tended to display peak recruitment toward the end of the retraction phase. Therefore, the specific effect of OC on the retraction module may be explained by the shorter interval between activity and reward compared to neurons in the protraction module.

Previous studies in vertebrates and invertebrates have similarly found that neuronal activity patterns underlying behavior lie within a low-dimensional subspace of the population space^1–3^. However, identifying a low-dimensional signature of OC has been challenging, in part because the number of cells from which one can record is commonly orders of magnitude smaller than the relevant population space. The isolated ganglia preparation allowed recording from a large proportion of the population, while removing variability due to sensory stimuli—the only stimulus was the reward. Therefore, we observed the direct effect of the association between fictive behavior and reward on the dynamics of the relevant neuronal population.

One possibility is that a learning signature could be found in changes to the low-dimensional subspace itself. However, primate studies suggest this subspace is resistant to change and imposes important constraints on learning-induced changes^5,24,25^. Accordingly, we found that neuronal activity before and after learning was consistently captured by a low-dimensional subspace, comprising protraction and retraction modules. The learning signature was not a change in the subspace, but rather in the relative *timing* of the trajectory within the subspace. Given the importance of timing actions for behavior production, learning may commonly act primarily by modifying neuronal module timing. Our findings can help guide characterization of learning signatures in systems with many more degrees of freedom, for which it may only be possible to record a small fraction of the relevant neuronal population.

## Methods

### Experimental subjects and preparations

*Aplysia californica* were obtained either from the National Resource for *Aplysia* of the University of Miami (Miami, FL) or from South Coast Bio-Marine LLC (San Pedro, CA). Animals (25–50 g) were kept in aerated artificial seawater tanks, and fed seaweed three times a week. Prior to each experiment, animals were food deprived for 3–5 days. A small piece of seaweed was used to test the animals’ motivation and feeding behavior prior to dissection. Animals were injected with a volume of isotonic MgCl_2_ equal in milliliters to 50% of the animals’ mass in grams. The buccal mass with buccal ganglia attached was isolated from the animal and placed in a Sylgard-coated chamber containing artificial seawater with a high concentration of divalent cations (HiDi-ASW; 210 mM NaCl, 10 mM KCl, 33 mM CaCl_2_, 145 mM MgCl_2_, 20 mM MgSO_4_, 10 mM HEPES, pH adjusted to 7.5 with NaOH). HiDi-ASW was used to reduce activity and muscle contractions during dissection. Buccal ganglia with buccal nerves were excised from the buccal mass and pinned down. The sheath of connective tissue around the ganglia was kept intact. Bipolar suction electrodes for recording and stimulation were positioned on each nerve (see below). HiDi-ASW was substituted with normal artificial seawater (ASW; 450 mM NaCl, 10 mM KCl, 10 mM CaCl_2_, 30 mM MgCl_2_, 20 mM MgSO_4_, 10 mM HEPES, pH adjusted to 7.5 with NaOH) prior to recording. Preparations were maintained at 15 °C throughout the experiment.

### Voltage-sensitive dye (VSD) imaging

Isolated buccal ganglia were stained for 7 min with 0.2 mg/ml of the absorbance VSD RH-155 (TRC Canada) in ASW. After staining, the solution was swapped for a lower concentration of RH-155 in ASW (0.01 mg/ml), which was kept for the remainder of the experiment. Using a Gilson Minipuls 3 peristaltic pump, the solution was continuously cycled through a Warner Instruments SC-20 inline cooler/heater under control of a Warner Instruments CL-100 temperature controller, which maintained the preparation at 15 °C. Imaging data was acquired by a Deep Well NeuroCMOS 128×128 camera (RedShirtImaging, SciMeasure) sampled at 1 kHz. The camera was fitted to an Olympus BX50WI microscope equipped with an Olympus 20× 0.95 NA XLUMPlanFL water immersion objective lens. The preparation was illuminated with a 150 W, 24 V Osram halogen light bulb powered by a Kepco ATE 75-8M power supply. Before reaching the preparation, light passed through a Brightline 710/40 nm bandpass filter.

Most neurons in the buccal ganglia are bilaterally symmetric, somata are located at the surface of the ganglia, and more neurons can be observed caudally than rostrally. Therefore, the number of recorded cells was maximized by imaging the caudal surface of the left hemiganglion, which accounts for about a quarter of the total surface area of the buccal ganglia. Each preparation was imaged for two 120 s periods, one before and one after either contingent or yoked training (see below).

### Extracellular electrophysiology

In a typical ingestion, the animal’s food-grasping radula is first protracted while open, and subsequently retracted back while closed around food. This biphasic protraction-retraction sequence can also be observed in the activity of nerves that control the feeding apparatus, both in intact animals^26^ and isolated ganglia^16,27^. Bipolar suction electrodes were used to monitor nerve activity associated with buccal motor patterns (BMPs). Nerve signals were amplified by A-M Systems model 1700 differential AC amplifiers, and digitized by Axon Instruments Digidata 1322A at 20 kHz. The following nerves were recorded: the left and right buccal nerves 2 and 3 (Bn2 and Bn3); right buccal nerve 1 (Bn1); left radula nerve (Rn); and left esophageal nerve 2 (En2). Electrical stimulation of En2, which acts as reinforcement, consisted of a 6 s train of 0.5 ms pulses at 10 Hz. Stimulation was manually triggered by the experimenter using a WPI Pulsemaster A300, and delivered by a WPI 850A stimulus isolator. Stimulus intensity was calibrated for each preparation by gradually increasing the intensity of a lower rate tonic stimulus (2 Hz, 0.5 ms) until a complete BMP was elicited. The efficacy of the regular 6 s stimulus was then confirmed after a brief rest. This procedure was devised to obtain a physiological threshold for En2 stimulation while minimizing the amount of stimulation prior to training.

### *In vitro* operant conditioning

After the En2 threshold was determined, experiments followed the timeline depicted in Fig. 1c. Preparations were allowed to rest for 40 min. Preparations that did not emit any spontaneous BMPs during this rest period were allowed to rest for an additional 20 min. The pretest period of 30 min was then initiated. To avoid a potential ceiling effect, and to ensure that preparations had a baseline rate of BMPs that was sufficient for training, preparations were discarded if they displayed <3 or >10 BMPs during the first two thirds of the pretest. All criteria were identical across groups. During the last third of the pretest, preparations were imaged for 120 s. If during this time no BMPs occurred, additional attempts were made to image during BMP generation. If no BMP could be captured before the end of the pretest, the preparation was discarded. After the pretest, a 30 min training period ensued. For contingent preparations, occurrence of an iBMP was followed by stimulation of En2. Any BMP elicited by En2 stimulation (i.e., initiated within 20 s after the offset of the stimuli train) was not reinforced. For each yoked preparation, En2 stimuli were identical to those received by its paired contingent preparation, irrespective of the BMPs emitted by the yoked preparation itself. Thus, yoked preparations controlled for any time-dependent or nonassociative effects. Preparations that had <3 effective En2 stimuli during training were discarded. An En2 stimulus was considered effective (for either contingent or yoked preparations) only if it clearly elicited at least one BMP. Upon conclusion of training, a 30 min posttest was performed. As in the pretest, preparations were imaged for 120 s during the last third of the posttest.

### BMP identification

During training, BMPs were manually classified by the experimenter in real time. For data analysis, BMPs were automatically identified and classified using custom MATLAB code. In both cases, the same criteria were used. Activity in Bn1 was used as an indicator of the timing of each BMP phase. Bn1 is particularly informative because it is active during protraction, inactive during retraction, and active again upon retraction termination. A BMP was identified when three criteria were met: (1) a burst of large-unit activity lasting at least 3 s occurred in Bn1; (2) the Bn1 burst was followed by a burst of large-unit activity in at least one of the two Bn3s; (3) at least one burst of large-unit Rn activity occurred during either the Bn1 burst (protraction) or the subsequent Bn1 suppression (retraction). Large-unit activity was defined as all activity with amplitude above a voltage threshold for each nerve, which was set independently for each experiment. The threshold for Bn1 was set to include all units that were consistently active during protraction and inactive during retraction. The Bn3 threshold was set to include only the largest units, which correspond to B4, a neuron known to be active at the onset of retraction. Similar to previous studies^28,29^, the Rn threshold was set to include units larger than baseline activity (i.e., activity occurring outside of BMPs). Thresholds were adjusted as necessary to ensure that the algorithm captured all BMPs occurring during the pretest, training, and posttest recordings for each preparation. Bursts were defined as at least 3 spikes with interspike intervals ≤4 s. BMPs were classified as ingestion-like (i.e., iBMPs) if ≥50% of the Rn burst duration occurred during the retraction phase^16,28^.

### Raw VSD data processing

VSD data were processed in a similar manner to that described elsewhere^15^. Briefly, regions of interest were drawn around the soma of each neuron, and the light intensity was averaged across pixels within the region for each time point. Traces for each cell were taken as the change in light intensity relative to baseline (i.e., ΔI/I), where baseline was defined as an average of the first 10 frames after shutter opening. Traces were filtered using MATLAB’s elliptic filter (n = 2, Rp = 0.1, Rs = 40, Wp = [0.02 0.2]).

### Denoising

Principal component analysis (PCA) was used to isolate and remove correlated noise, such as that associated with movement. PCA extracts a set of components from the data that can be ordered by the amount of variance-covariance in the data explained by each component. If some components consist primarily of noise, the data can be reconstructed without those components to improve signal-to-noise ratio. At the millisecond timescale, most of the correlation between the traces of individual neurons was due to noise that affected many neurons simultaneously, such as tissue movement, vibrations, and fluctuations in illumination. This was the case because the amplitude of RH-155 signals is very small (i.e., in the order of 1×10^−4^ ΔI/I), and, importantly, because action potentials in *Aplysia* are not correlated at the millisecond scale during spontaneous BMP generation. Note that this is different from the correlation between firing rates examined throughout this paper, which is on the scale of seconds. Thus, we performed PCA on the z-scored traces from all neurons in a preparation, removed the top *n* components accounting for most of the correlation between traces, and reconstructed the data. The number of components to remove was determined by performing reconstructions with *n* ranging from 0 to all components, and selecting the most de-correlated reconstruction. We validated this procedure by adding artificial action potentials of varying amplitudes to a test data set, performing the full range of reconstructions, and attempting to detect the artificially added action potentials using the algorithm described below. The test data were a set of 100 traces from a real preparation. Because a substantial proportion of the added spikes were buried in real noise, they served as a realistic test of signal improvement by denoising. Consistent with visual inspection of signal quality, the most de-correlated reconstructions led to the highest true positive rates.

### Spike detection

Spikes were detected in a neuron when a downward deflection in the VSD trace exceeded five times an estimate of the standard deviation of the noise. This estimate was given by

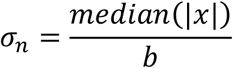

where *x* is the trace of the neuron, and *b* = 0.6745 is the constant for the relationship between the median absolute deviation estimator and the standard deviation *σ*_*n*_ for Gaussian distributions^30–32^. A re-arm window of 10 ms was used to prevent a single action potential from being counted more than once. This simple threshold method yielded comparable true positive rates as did a more sophisticated spike detection algorithm^33^ in a test data set to which artificial action potentials of varying amplitudes were added.

### Non-negative matrix factorization (NNMF)

NNMF is a dimensionality reduction approach that optimizes an approximation of the data matrix *X* from a pre-determined number of components *k*, given non-negativity constraints (for a detailed overview, see ^17^). NNMF is well-suited to identify a low-dimensional signature of learning in the *Aplysia* feeding circuit for at least three reasons. First, NNMF has been used to identify groups of neurons participating in memory^34^. Second, NNMF has successfully decomposed behaviors into their motor module building blocks^18^. Third, the assumptions and constraints adopted by NNMF are compatible with neuronal activity data^17^. For the activity of a neuronal population over time, NNMF can be described by

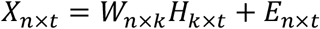

where *X*_*n*×*t*_ is the activity of *n* neurons over *t* time points, *W*_*n*×*k*_ is a matrix of the contributions of each of the *n* neurons to each of the *k* components, *H*_*k*×*t*_ is a matrix of the magnitude of each of the *k* components at each time point *t*, and *E*_*n*×*t*_ is the approximation error or residual term that NNMF minimizes. Note that these *k* components are referred to as modules throughout this manuscript.

NNMF was performed using the seqNMF MATLAB package developed by Mackevicius et al.^35^ with L = 1, λ = 0, and otherwise default parameters. This set of parameters corresponds to standard NNMF using multiplicative update rules first introduced by Lee and Seung^37^. Six repetitions were performed for each run to ensure that the algorithm consistently converged to the same modules. Given consistent convergence, an arbitrary repetition was ultimately selected.

Prior to NNMF, data were temporally smoothed by convolving the spike train of each neuron with a Gaussian. Because a difference in pairwise correlations was observed when spikes were binned with 1 s bins (Fig. 2d), the Gaussian was set to have a standard deviation of 1 s. This smoothing is compatible with the duration of BMPs as well as *in vivo* feeding behaviors, which usually unfold over the course of tens of seconds. After smoothing, pretest and posttest data were concatenated. This held the contribution matrix *W* constant between pretest and posttest, allowing for definition of identical components (i.e., modules) before and after training. Thus, we were able to compare the time course of recruitment of the same modules before and after training.

Given that overall BMP duration and the duration of each phase vary from pattern to pattern, comparing component recruitment during BMP generation required normalizing these durations. This normalization was performed by using linear interpolation to resample component recruitment during each phase of each BMP to a standard of 180 time points. Finally, this resampled time course of module recruitment was averaged across BMPs to obtain a mean pretest and a mean posttest time course for each preparation. The timing and magnitude of the peaks of these average time courses were then compared (Fig. 3d–g). Furthermore, the average time courses during the pretest (i.e., before any training) were used to examine the consistency of modules across preparations (Extended Data Fig. 1).

A critical factor in NNMF and other dimensionality reduction techniques is selection of the number of modules *k* used to approximate the data. Various strategies have been used to make this selection^6,17,34^. Examples include using heuristic solutions (such as a simple threshold on the approximation error or variance-covariance explained by the reconstruction), or using estimates of the tradeoff between information loss and model complexity (e.g., Akaike’s or Bayesian information criteria), among others. Here, priority was given to ensuring that modules were behaviorally relevant and consistent across animals, thereby allowing for direct comparisons of the effects of learning on module recruitment. We assessed this cross-animal consistency by computing the correlation between the average pretest time course of module recruitment during a normalized BMP across preparations (Extended Data Fig. 1a–b). We performed a series of factorizations with *k* ranging from one to ten, each repeated six times. For each repetition, a matching algorithm sorted modules into *k* clusters by matching modules in the order of their correlations across preparations. Within preparation matches were disallowed, and each module was only allowed to match with one other module per preparation. Note that this is distinct from the hierarchical clustering in Extended Data Fig. 1b, which has no such limitations. This matching algorithm ensured that each preparation had exactly one module in each of the *k* clusters. An overall mean correlation among matched modules was computed for each NNMF run as the final consistency metric (Extended Data Fig. 1c).

The number of modules *k* = 2 was selected because it yielded the most consistent modules across preparations, and because it did so while explaining most (74.9 ± 1.8%) of the total power in the data. Explained power is a metric of how well a factorization approximates the original data, analogous to the concept of explained variance-covariance in PCA. The explained power was computed as in Mackevicius et al.^36^, that is

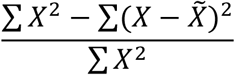

where *X* is the original data, 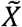 is the data reconstruction, and ∑ *X*^2^ denotes 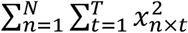.

Hierarchical clustering (Extended Data Fig. 1b) was performed with MATLAB’s *linkage* function using the *average* method. Briefly, the function starts with single-module clusters and progressively forms larger clusters by grouping the most similar (i.e., most correlated) clusters, until a single cluster encompasses all modules. Once a module is assigned to a cluster, the module is no longer individually compared to others; rather, a comparison is made between the entire clusters. The dissimilarity between two clusters is equal to the mean dissimilarity among all possible pairs of modules between the two clusters.

### Spatial distribution of motor modules

To allow for comparisons of module localization across animals, all preparations were aligned to a reference ganglion (preparation A in Extended Data Fig. 2a). A regression of the pixel coordinates of the largest neurons (i.e., the upper quartile for size) was used to approximate the orientation and offset of each ganglion^15^.

### Statistical hypothesis testing

All hypothesis tests were performed on SigmaPlot 12. Normality was not assumed and two-tailed Wilcoxon signed-rank tests were used to perform paired comparisons in Figs. 1f–g, 2d–e and 3f–g. Alpha was set to 0.05. Data analyses were performed while blinded to the training history of each preparation.

## Acknowledgments

We thank Drs. Ryota Homma and Paul Smolen for comments on an earlier draft of the manuscript.

## Author contributions

Conceptualization: RMC, DAB, JHB

Methodology: RMC, DAB, JHB Investigation: RMC

Visualization: RMC Analysis: RMC

Funding acquisition: JHB

Project administration: JHB

Supervision: DAB, JHB

Writing – original draft: RMC

Writing – review & editing: RMC, DAB, JHB

