## Supplementary Figures for "Neuronal population activity dynamics reveal a low-dimensional signature of operant learning"

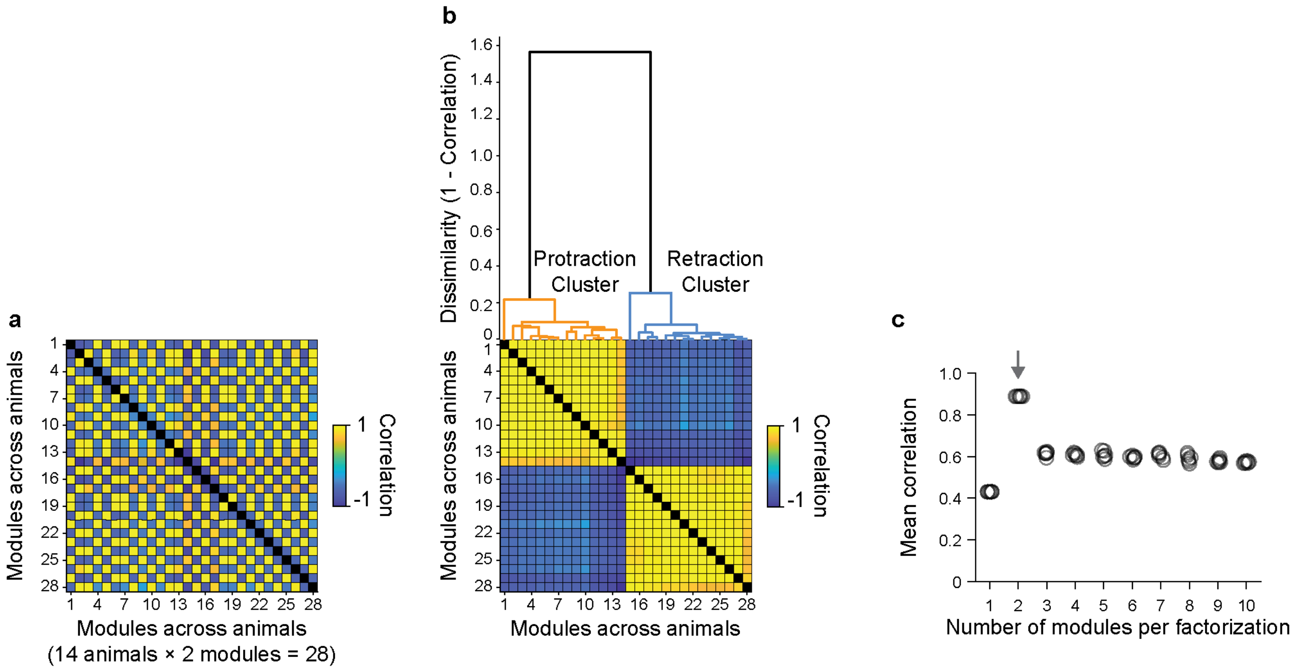


**Extended Data Fig. 1 | Consistency of modules across preparations. a**, Matrix of the correlations between the time course of recruitment of all pairs of modules across preparations during a normalized BMP. Two modules were obtained by NNMF per preparation. Correlations were computed using pretest data alone. Modules are sorted by preparation. Self-correlation diagonal is blacked out for clarity. **b**, Hierarchical clustering of modules based on correlations between their recruitment time courses. Dendrogram (top) reflects clustering by average group linkage (see text for details). Correlation matrix (bottom) is the same as panel a, but sorted by cluster affinity. Each preparation had one module in the protraction cluster, and one in the retraction cluster. **c**, Overall correlation among matched modules across preparations for various NNMF runs with a range of numbers of modules (see text for matching algorithm). Circles represent individual NNMF runs. Arrow indicates the number of modules selected for analyses throughout the paper.

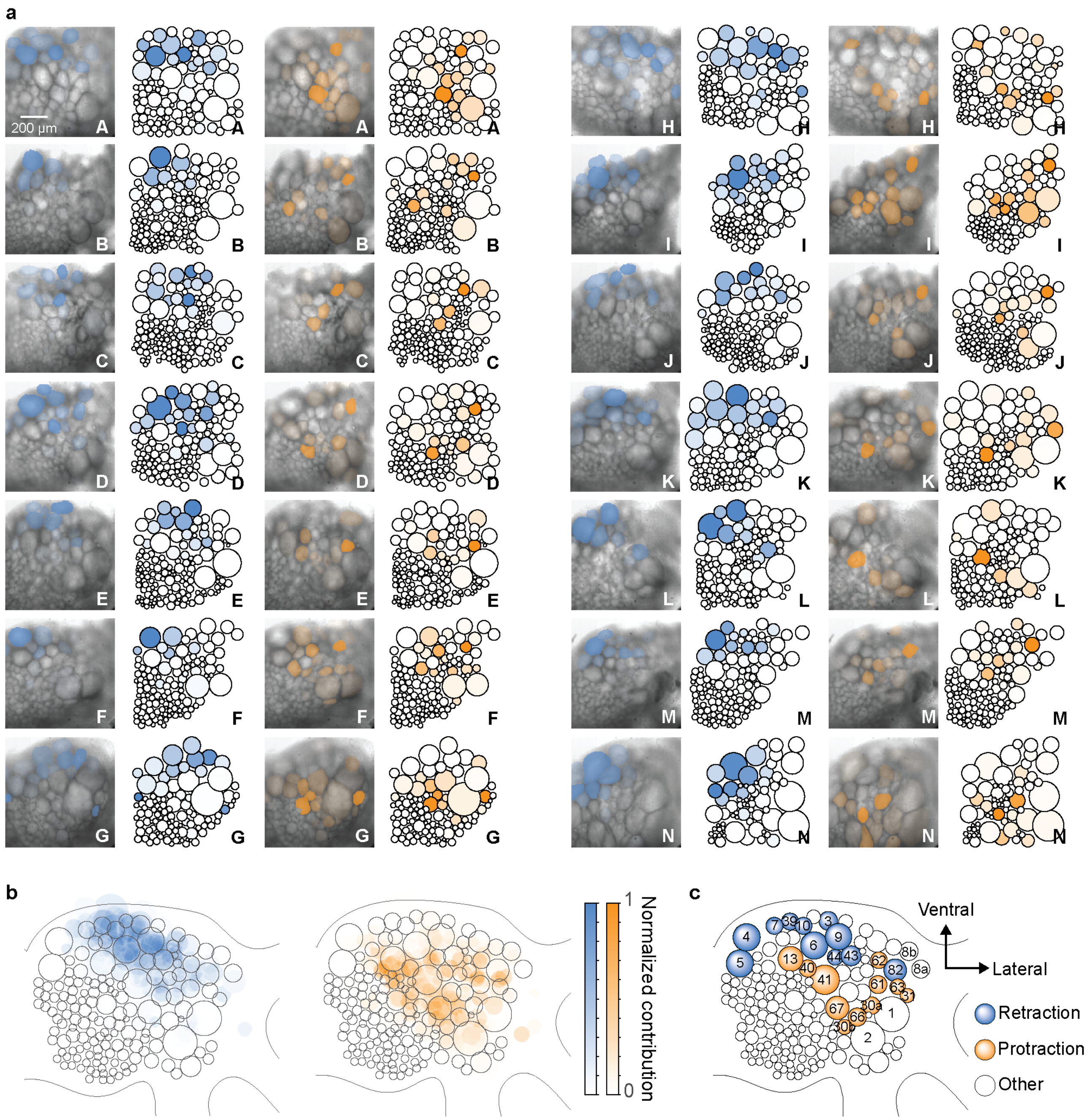


**Extended Data Fig. 2 | Spatial distribution of neurons contributing to motor modules. a**, Contributions made to the modules by each recorded neuron across all preparations. Preparations are identified as A–N. Colors indicate the cell's contribution to the retraction (blue) or protraction (orange) module. Each image is accompanied by a diagram displaying neuron sizes, positions, and module contributions. Color scales are as in panel b. **b**, Overall localization of neurons contributing to the retraction (left) and protraction (right) modules across preparations. Ganglia, and diagrams, are aligned to preparation A in panel a, and the color intensity is averaged across preparations. **c**, Localization of neurons characterized in the literature that are active during retraction or protraction phases.
